# Gaussian curvature and the budding kinetics of enveloped viruses

**DOI:** 10.1101/457135

**Authors:** Sanjay Dharmavaram, Baochen She, Guillermo Lázaro, Michael F. Hagan, Robijn Bruinsma

## Abstract

The formation of a membrane-enveloped virus such as HIV-1 starts with the assembly of a curved layer of capsid proteins lining the interior of the plasma membrane (PM) of the host cell. This layer grows into a spherical shell enveloped by a lipid membrane that is connected to the PM via a curved neck (“budding”). For many enveloped viruses the scission of this neck is not spontaneous. Instead, the elaborate “ESCRT” cell machinery needs to be recruited to carry out that task. It is not clear why this is necessary since scission is spontaneous for much simpler systems, such as vesiculation driven by phase-separation inside lipid bilayers. Recently, Brownian dynamics simulations of enveloped virus budding reproduced protracted pausing and stalling after formation of the neck [1], which suggest that the origin of pausing/stalling is to be found in the physics of the budding process. Here, we show that the pausing/stalling observed in the simulations can be understood as a purely *kinetic* phenomenon associated with a “geometrical” energy barrier that must be overcome by capsid proteins diffusing along the membrane prior to incorporation into the viral capsid. This geometrical energy barrier is generated by the conflict between the positive Gauss curvature of the capsid and the large negative Gauss curvature of the neck region. The theory is compared with the Brownian simulations of the budding of enveloped viruses.

**Author summary:** Despite intense study, the life-cycle of the HIV-1 virus continues to pose mysteries. One of these concerns the assembly of the HIV-1 virus inside infected host cells: it is *interrupted* at the very last moment. During the subsequent pause, HIV-1 recruits a complex cell machinery, the so-called “ESCRT pathway”. The ESCRT proteins pinch-off the “viral bud” from the host cell. In this paper, we propose that the reason for the stalling emerges from the fundamental physics of the lipid membrane that surrounds the virus. The membrane mostly follows the spherical geometry of the virus, but in the pinch-off region the geometry is radically different: it resembles a neck. By combining numerical and analytical methods, we demonstrate that a neck geometry creates a *barrier* to protein entry, thus blocking proteins required to complete viral assembly. This “geometrical barrier” mechanism is general: such a barrier should form during assembly of *all* membrane-enveloped viruses – including the Ebola and Herpes viruses. Indeed many families of enveloped viruses also recruit the ESCRT machinery for pinch-off. A fundamental understanding of the budding process could enable a new strategy to combat enveloped viruses, based on selective stabilization of membrane neck geometries.

## Introduction

Many viruses that infect animals, including many human pathogens, are surrounded by a lipid membrane that allows the virus to enter a host cell by membrane fusion [2]. The envelope also may help to avoid attack by the immune system. Well-known examples are the retroviruses, like HIV-1, the Herpesviruses, and the Filoviruses (e.g. Ebola virus). Apart from some viral proteins, the viral membrane is derived by *budding* of the virus outer protein shell (capsid) through a lipid-based membrane of its infected host cell, such as the plasma membrane (PM). In many enveloped virus families, budding is driven by the assembly of a curved layer of the viral capsid proteins that lines the interior of the PM the host cell [3–5]. For single-stranded RNA viruses like HIV-1, viral RNA genome molecules are associated with the proteins on the interior side of the layer [6]. This layer grows, by transport of proteins from the cytosol where the capsid proteins are being synthesized, and evolves into a spherical-cap shape covered by a membrane still connected to the PM via a curved neck. Figure 1 shows a sketch of a typical late-stage bud of an HIV-1 viral particle (“virion”) as obtained from cryo-EM tomograms [3].

**Fig 1.**
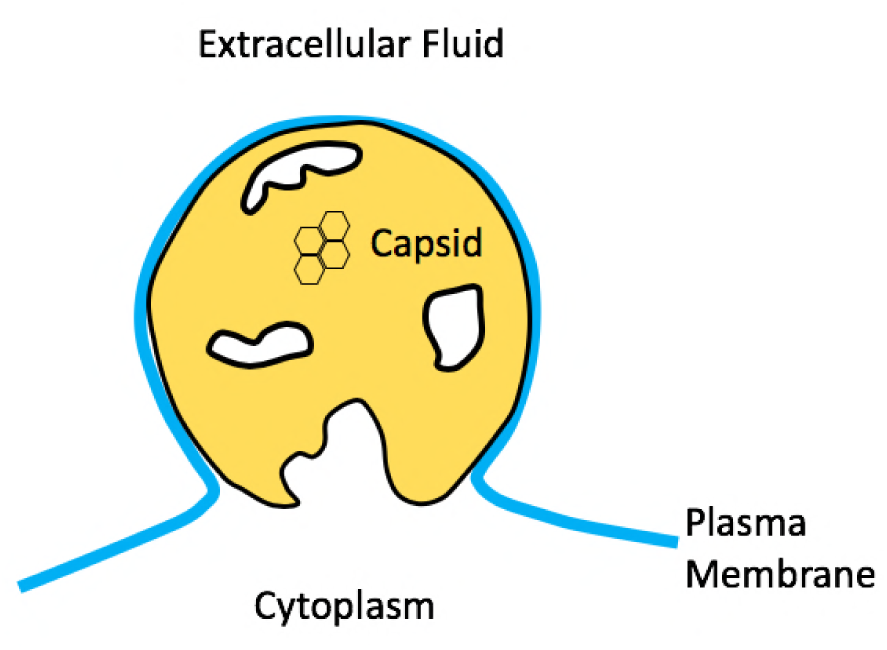
Sketch of a budding HIV-1 virion just prior to pinch-off. The lipid bilayer covering the capsid is connected to the plasma membrane by a highly curved neck. The capsid proteins mostly are arranged in hexons but the capsid has holes. The large hole surrounding the pinch-off site remains in the completed virion.

Up to that point, budding is a spontaneous process driven by attractive interactions between the capsid proteins. For many – but not all – enveloped viruses the final scission of the membrane neck requires ”hijacking” the *ESCRT* machinery of the cell [7, 8]. ESCRT is a complex of proteins involved in cellular processes that require membrane scission, such as the formation of multi-vesicular bodies and cytokinesis [9]. Scission may be able to proceed without ESCRT. For example, scission of HIV-1 buds does take place in the absence of the ESCRT machinery but there is a significant delay [10, 11]. It seems likely that ESCRT recruitment is necessary to assure that scission takes place *on time.* For the example of HIV-1, scission has to take place before activation of the autocatalytic protease process that disintegrates the capsid proteins [12]. The fact that ESCRT recruitment takes place across so many different families of enveloped viruses suggests that the origin of the pausing/halting kinetics must be found among basic properties of the budding process shared among all enveloped viruses. No cell-biological process has been identified so far. In this paper we propose that a physical mechanism is responsible for the pausing/stalling.

## Results and Discussion

Figure 1 showing an HIV-1 bud provides us with some clues. The lattice of capsid proteins of an HIV-1 bud has a large *hole* surrounding the pinch-off site. The boundary of the hole represents a linear interface between the part of the PM that is covered by capsid proteins and the part that is not. This hole survives the pinch-off process and only about 2/3 of the membrane of the completed immature HIV-1 virus is covered by proteins. It is important to note that hole formation is not observed for other enveloped viruses such as the Herpes and alphaviruses. Figure 1 suggests that the transport current supplying proteins to the curved neck region somehow has “dried-up” before the spontaneous part of the assembly could complete. This conclusion is supported by numerical simulations. Figure 2 shows snapshots of a Brownian Dynamics simulation of a simple, coarse-grained model of the budding of the enveloped protein shell of the alphavirus (composed of transmembrane glycoproteins (GPs) [1]. The GPs were modeled as rigid trimers of truncated cones, with each cone comprising a linear array of six beads of increasing diameter. The cone angle was set so that in the absence of a membrane, the GPs assembled into hollow, roughly icosahedral shells containing 80 trimers, though they form larger shells in the presence of a membrane [1]. The membrane was represented by the implicit solvent model of Cooke and Deserno [13]. As the assembly proceeded, the aggregate of GPs adopted the shape of a *spherical cap* that gradually closed. For high values of the protein-protein binding energy *ϵ*_gg_, complete closure was achieved and pinch-off was *spontaneous*. The resulting spherical shells were highly defected. The growth rate was non-uniform: the assembly rate started to slow down when the shells reached approximately 2/3 completion. Slow-down became more pronounced with decreasing *ϵ*_gg_ while the final spherical shells were less defected. A critical value was reached for *ϵ*_gg_ about 1.7*k*_B_*T*. Below this value, the assembly process stalled before closure could be achieved. The diameter of the remaining neck grew larger as *ϵ*_gg_ was further decreased. Figures 1 and 2 suggest a physical mechanism for the stalling: the part of the membrane linking the bud to the PM has a very different geometry from the part of the membrane covering the proteins shell: the latter has a modest spherical curvature, while the neck as a highly curved hyperbolic shape. Could the energy cost of the bending the PM membrane in the neck region generate a barrier for the scission process?

**Fig 2.**
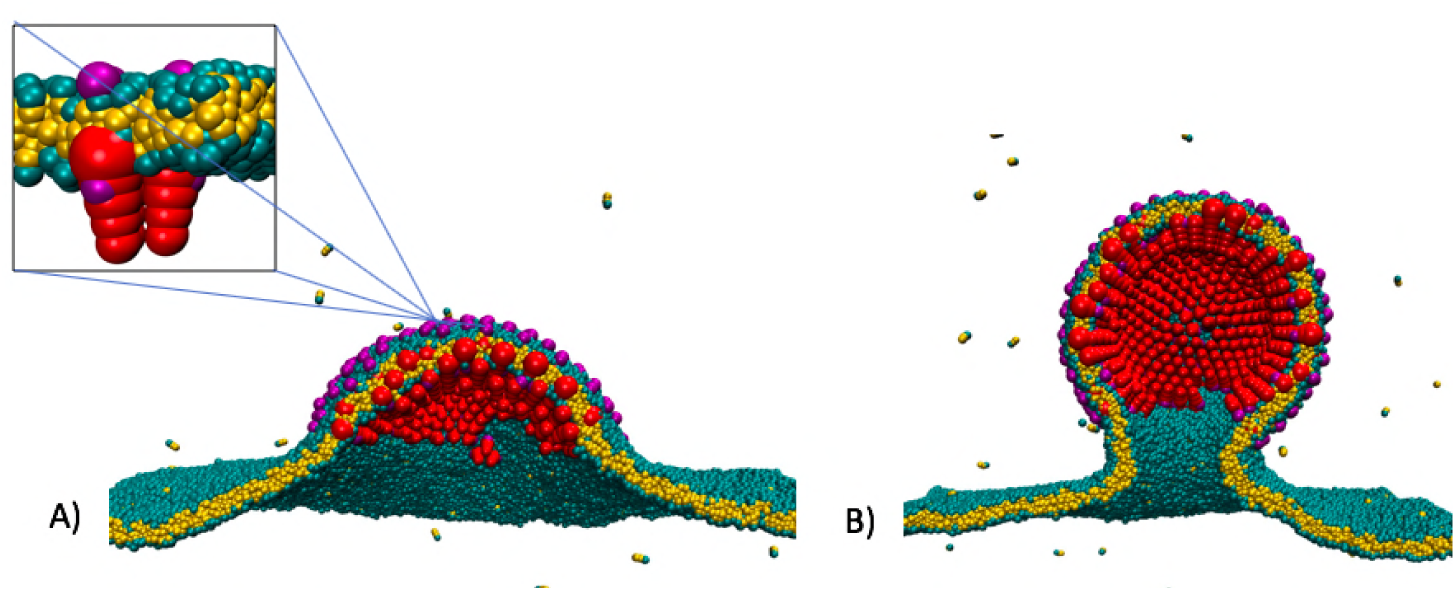
Brownian Dynamics simulation of the budding of an enveloped virus using a coarse-grained model of the lipid molecules and capsid proteins of the alpha virus (square inset). The strength *ϵ*_gg_ of the interaction between the capsid proteins was 2*k_B_T*. A) Snapshot of the simulation at an early stage of the bud. B) Snapshot of the bud during pausing.

To investigate the physics of such a scenario, start with the simple case of a *flat* circular disk of assembled capsid proteins. For now, assume that the disk is not attached to a lipid bilayer. If the radius of the disk is **ρ** then the assembly free energy *F*_0_(**ρ**) equals π**ρ**^2^Δ*μ*+2π*ρτ*. In the first (area) term, Δ*μ* represents the difference between the chemical potential per unit area of unbound capsid proteins and capsid proteins that are part of the disk. In the second (perimeter) term, *τ* is the difference in chemical potential per unit length of capsid proteins on the perimeter of the disk and of capsid proteins in the interior. For the capsid proteins of small viruses, Δ*μ* has been measured to be of the order of 5 – 10 *k_B_T* per square nanometer [14] though for the highly defected capsid of HIV-1 it is probably less. It is also known that the critical nucleus for assembly of HIV-1 is of the order of just a few capsid proteins [3]. It follows that the radius *τ*/**ρ** of the critical nucleus must be of the order of a few nanometers, meaning that *τ* must be of the order of a few *k_B_T* per nanometer. Now, allow the disk to curve out of the plane. The bending energy of the disk will be described by the Helfrich bending free energy for deformable surfaces, which has been used to elucidate the physics of lipid bilayers [15] and viral capsids [16]. It also can account for the spontaneous budding of multi-component lipid bilayers [17–19]. The Helfrich bending energy *F_B_* of a surface is defined as

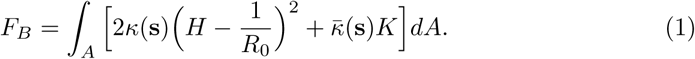

In the first term, 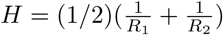 is the mean curvature with *R*_1,2_ the principle curvature radii at a point **s** on the surface of the disk, while 1/*R*_0_ is the preferred or spontaneous curvature of the protein layer. The coefficient *κ*(*s*), the local bending modulus, is positive. The second term is the Gaussian curvature energy with *K* = 1/(*R*_1_*R*_2_) the Gaussian curvature. *K* is positive for a spherical surface, such as a spherical virus, and negative for a hyperbolic surface (such as the neck). The Gaussian curvature modulus 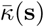 in principle can have either sign but for the present case it should be negative. For now, assume that the two moduli are constant. When the bending energy *F_B_* is added to the assembly free *F*_0_(**ρ**) of the disk then minimization of the total free energy with respect to shape produces a curved shell with the geometry of a spherical cap if 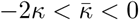. The curvature radius of the spherical cap is 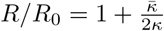. However, in the limit that 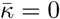, *any* surface of constant mean curvature obeying 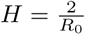 represents a minimum energy state. Spherical caps obey this condition but so do cylinders, unduloids and tubular structures. The presence of a negative Gaussian curvature modulus is thus a requirement for the stability of a protein shell in the shape of a spherical cap.

The Gaussian curvature energy has not entered in discussions of the budding of lipid bilayers nor in previous models of the budding of viruses [20–22] and clathrin cages [23] because of the Gauss-Bonnet Theorem (GBT). According to the GBT, the integral of the Gaussian curvature *K* over a surface A with boundary S obeys 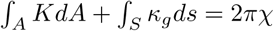 with *κ_g_* the *geodesic curvature* of the boundary and with *χ* a topological invariant (know as the “Euler characteristic”). For a closed membrane without boundary that has a uniform Gaussian curvature modulus, the area integral of the Gauss curvature equals 4*π*. If the topology of the surface does not change then the Gaussian curvature energy plays no role. This is not true if the Gaussian modulus is not a constant. Consider a closed membrane composed of two different parts that are joined along a closed boundary line *S.* One part has moduli *κ*(*s*) = *κ_C_* and 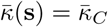 while the other part has moduli *κ*(*s*) = *κ_L_* and 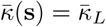. Application of the GBT now leads to

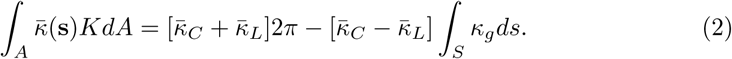

For such a non-uniform surface, the Gaussian curvature energy 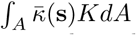 is dependent on the geometry of the boundary line and no longer a topological invariant.

Now assume that the protein layer adheres to a lipid bilayer. Consider the geometry shown in Fig. 3, which represents a bud prior to pinch-off. A spherical-cap – representing the partially assembled capsid – is attached to a minimal surface in the form of a *catenoid of revolution* –representing the protein-free lipid bilayer (both shapes are extrema of FB). The interfacial boundary between them is assumed circular. The apex angle of the cone subtended by the center of the sphere and the boundary line is denoted by *α,* the “closure angle”. The area of the spherical cap must equal that of the flat disk before bending (*π*ρ**^2^), which leads to the geometrical relation **ρ** = 2*R* cos *α*/2. The complete continuum energy *F* = *F*_0_ + *F_B_* is

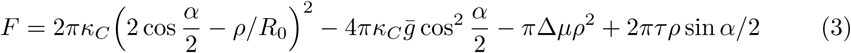

**Fig 3.**
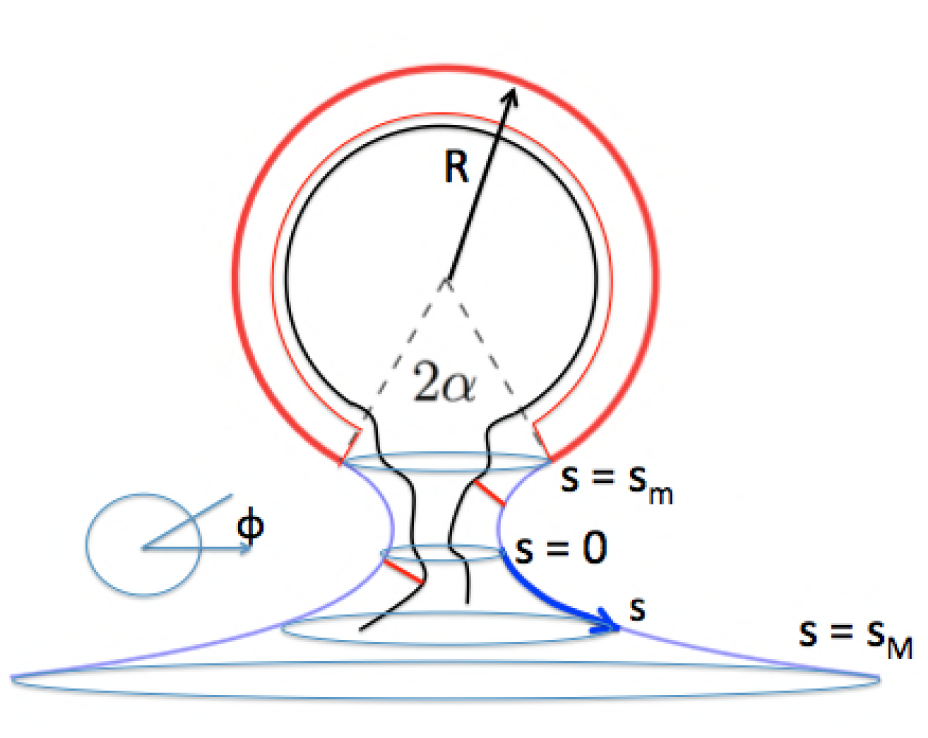
Schematic geometry of a late-stage bud prior to pinch-off. Blue line: bare lipid bilayer membrane. Heavy red line: lipid bilayer attached to a curved layer of capsid proteins shown as a thin red line. Black line: RNA genome molecules associated with the bud.The boundary between the two bilayers and the center of the sphere spans a cone with aperture angle 2*α*. Two monomeric capsid proteins diffusing along the lipid bilayer are shown as two red bars associated both with the membrane and an RNA genome molecule. In the curvilinear coordinate system (*s*,*φ*), *s* measures the shortest arc distance between a point and the cross-section with minimum diameter (s=0). It ranges from *s_M_ >* 0 to *s_m_ <* 0. Finally, *φ* is the azimuthal angle of the circle on the surface perpendicular to the central axis on which the point is located.

The first term is the first part of the bending energy *F_B_,* which involves the mean curvature. The second term is Gaussian curvature energy, computed using Eq.2. The dimensionless parameter 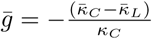 measures its strength. Note that 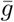 vanishes when the two layers have the same Gaussian curvature modulus. Since the moduli of the capsid are expected to be larger in magnitude than those of the lipid bilayer [24–27], it is expected that 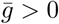.

If the capsid shell is best treated as a two-dimensional visco-elastic system – for example because the shell is highly defected – then *α* should be treated as a variational parameter to be determined by free energy minimization. The function *F*(*α*) has two extrema: one at a non-zero *α* = *α*^*^(**ρ**) that corresponds to an open spherical cap and one at *α* = 0 that corresponds to a closed shell. A typical example of *α*^*^(**ρ**) is shown in Fig.4. As **ρ** increases from zero, *α*^*^(**ρ**) decreases monotonically from *α*^*^(0) = *π.* Initially, the spherical cap state is the minimum free energy state, but at a point **ρ** = **ρ**^*^, the energy of the *a* = 0 closed shell state drops below that of the *α* = *α*^*^(**ρ**) spherical cap state. At that point, the spherical cap state is connected via a neck to the membrane. At a slightly larger value of **ρ*,* the spherical cap state becomes locally unstable and abruptly transforms at fixed **ρ** to a closed shell state. This scission step is driven by the line-tension term *τ.* There is no energy activation barrier: scission is spontaneous in continuum theory.

**Fig 4.**
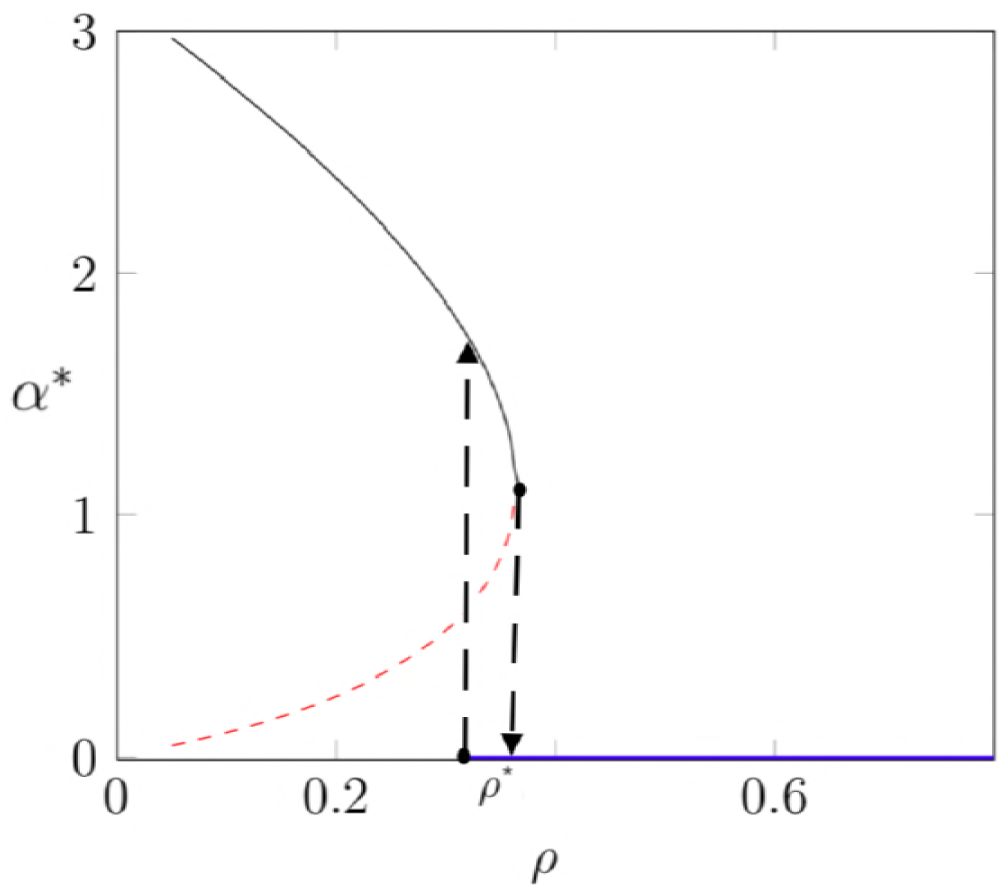
Example of the dependence of the aperture angle *α*^*^ that minimizes the free energy on the growth parameter **ρ*.* The partial shell in the shape of a spherical cap with *α*^*^ *>* 0 is locally stable along the black line and unstable along the red dashed line. The completed capsid with *α*^*^ = 0 is stable along the blue line. The black dashed lines mark limits of local stability. Parameter values: *κ_C_* = 0.5, Δ*µ* = 0, 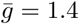, and *τ* = 0.5. The growth parameter **ρ** is expressed in units of *R*_0_ = 1.

If the capsid shell is better treated as a positionally ordered layer then an *elastic strain energy* must be added to the bending energy. Energy minimization of curved elastic shells is a more complex problem [28]. This is related to the mathematical necessity of introducing five-fold symmetry sites in an hexagonal lattice that covers a curved shell as it grows in size. This is necessary in order to reduce curvature-induced elastic stress [29, 30]. The minimum free energy state *E*(*n*) of complete shells is known to have a complex dependence on the number of particles *n* in the shell [31]. For a positionally ordered icosahedral capsid, *E*(*n*) is expected to have a sharp minimum at the particular *n* value that corresponds to that of the completed capsid. General statements can still be made about the way the introduction of elasticity affects shell closure because, for fundamental reasons, local changes of the Gaussian curvature of an elastic shell generate shear stress ^1^. The first statement is that the discontinuous transformation at fixed particle number of an open spherical cap into a closed sphere that we just described for a fluid shell is not possible for an elastic shell (unless the hole is very small): the closure of the shell is a continuous process. More generally, *if* the growing shell can be described as a spherical cap *then* the radius of curvature of the shell must be constant in time and hence equal to the curvature radius of the assembled shell. It follows that the relation between the curvature parameter *α* and the growth parameter **ρ** now is fixed by the condition **ρ** = 2*R_c_* cos *α/*2 with *R_c_* the capsid radius.

Brownian dynamics simulations of alphavirus budding produce capsids that are relatively ordered with a mean curvature that does not change significantly during budding. This indicates that for these simulations, the elastic shell description is more appropriate. Could curvature-induced elastic stress somehow be responsible for the pausing and stalling? This can be tested by repeating the simulations of Fig.2 in the absence of a lipid bilayer and with monomers in solution freely diffusing to the edge of the shell. Pausing and stalling effects disappeared completely while the curvature radius was similar. The removal of the lipid bilayer should not change the physics of curvature-induced elastic stresses inside the protein shell. We conclude that pausing/stalling is not related to curvature-induced elastic stresses.

Could stalling and pausing then be a purely *kinetic* problem associated with the transport of capsid proteins along the curved lipid bilayer to the growing shell? To investigate this possibility, allow for a low concentrationd Φ of capsid proteins diffusing in from infinity while adhering to the membrane *outside* the partially assembled shell. We will retain the geometry of Fig.3 with *R* = *R*_0_, and look for a steady-state diffusion current from infinity to the growth interface. Use the curvilinear coordinate system (*s,φ*) shown in Fig.3, where *s* measures the arc distance of a point on the surface to the minimum cross-section of the neck (at s=0), and where *φ* measures the azimuthal angle. Assuming rotational symmetry under steady state, the surface protein concentration Φ(*s*) in the neck region should only depend on *s.* Far from the neck area, the capsid protein concentration is set equal to a constant Φ_0_. We will assume that a capsid protein arriving at the growth interface is immediately absorbed into the shell, which means that Φ = 0 at the growth interface (“absorber” boundary condition). The diffusion current density *J* of a low concentration of particles on a curved surface [32] has the general form

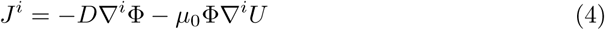

with *D* the surface diffusion coefficient, *µ0* = *D/k_B_T* the capsid protein mobility, *i* a component of the curvilinear coordinate system of Fig.3, and *U*(*s*) the potential energy of a capsid protein at point *s* on the surface of the curved lipid bilayer. Under steady-state conditions,

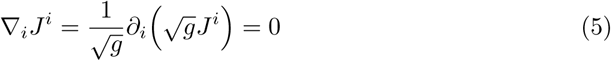

For a catenoid surface of revolution, the determinant *g* of the metric tensor obeys 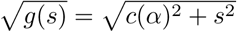 with *c*(*α*) = *R*_0_ sin^2^(*α*) the minimum radius of the neck. The current conservation equation then reduces to

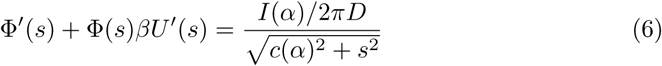

where *I*(*α*) is the total incoming current that we need to determine. This equation is solved by the Ansatz Φ(*s*) = *p*(*s*)*e^−βU^*^(*s*)^ where 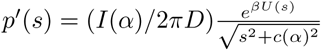. Impose absorber boundary conditions Φ(*s_m_*) = 0 along the growth interface at *s* = *s_m_* and set Φ(*s_M_*) = Φ_0_ far outside the neck area at *s* = *s_M_,* where we also place the zero of the potential energy (so *U*(*s_M_*) = 0). This gives for the current:

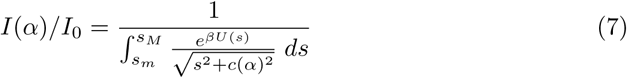

with *I*_0_ = 2*πD*Φ_0_. If the metric factor 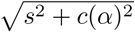 is set to one in Eq.7, then it reduces to a form similar to the Kramers expression for steady-state diffusion in a potential [33]. The non-entropic contribution to the chemical potential of a protein is given by 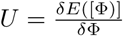 where E([Φ)] is the internal energy of the lipid bilayer outside the capsid but with a low concentration of proteins. By repeating the symmetry arguments that are used to obtain the Helfrich bending energy for a curved membrane [15], one finds that E([Φ)] must have the general form

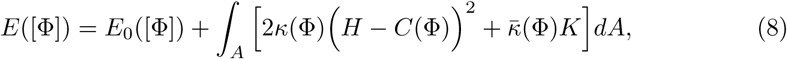

Here, *E*_0_([Φ]) is the protein-membrane interaction energy of a flat membrane while *κ*(Φ) and 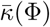 are the concentration-dependent curvature moduli. Finally, *C*(Φ) is the concentration-dependent spontaneous curvature. In the limit Φ = 0, all these quantities should reduce to the values appropriate for a pure lipid bilayer with capsid proteins. Expanding to lowest order in Φ around this state gives

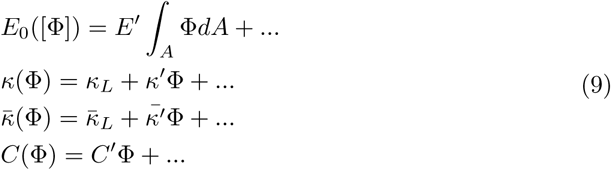

It follows that

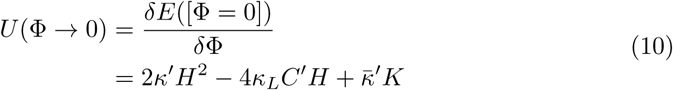

In the last step we assumed that at *s* = *s_M_,* where *U* = 0, the membrane is flat so *H* = *K* = 0, which means that 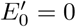. Finally, since *H* = 0 for a minimal surface, we arrive at the simple result that 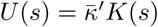 to first order in Φ. Since we saw earlier that 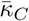 must be negative for a stable spherical cap state, we conclude that 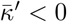. For a catenoid of revolution the Gaussian curvature is given by 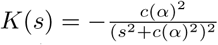 so 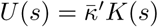 has a maximum at the center of the neck (*s* = 0). There is thus a “geometrical” energy barrier that the diffusing proteins need to overcome before they can be absorbed at the growth interface.

For reference, consider first the case where coupling to the Gaussian curvature is neglected. In that case, Eq.7 for *U* = 0 reduces to

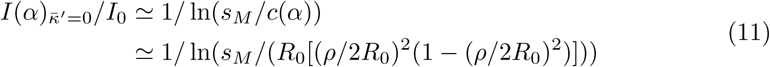

In the last step we used the fact that **ρ*/R*_0_ = 2 cos *α*/2. The appearance of a logarithmic dependence on the system size *s_M_* is typical of two-dimensional diffusion problems. The current first increases with **ρ** until the capsid has the shape of hemisphere at the point (**ρ**/2*R*_0_)^2^ = 1/2. Afterwards, the current decreases back to zero. A plot of the current *I* as a function of **ρ**^2^ is symmetrical around the midpoint maximum 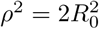.

Now consider the case that 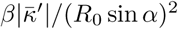 is comparable to, or larger than, one. Using the steepest descent method around the maximum of *U*(*s*) at *s* = 0 leads to

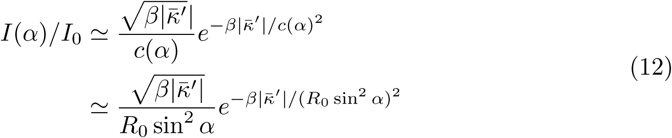

The current as a function of the aperture angle has an essential singularity at the pinch-off point *α* = 0 and **ρ*/R*_0_ = 2, where it precipitously drops to zero. A plot of the current as a function of **ρ**^2^ is highly asymmetric. Figure 5 compares Eq.7 with the outcome of the simulations of Fig.2. The only fitting parameter was the dimensionless strength *γ* of the coupling of the diffusing proteins to the Gaussian curvature of the membrane. While Eq. 7 is reasonably consistent with the data, the data scatter does not allow for a definitive quantitative comparison. It is however possible to make qualitative comparisons. Recall that the presence of a maximum in the current profile is a feature as well of conventional diffusive transport (i.e., *γ* = 0 as noted by a number of authors [34–37]). In the absence of the geometrical barrier, the maximum should be at 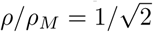 where the shell has the shape of a hemisphere (located by an arrow). The position of that maximum disagrees strongly with the maximum in data of Fig.5. We can definitely rule out conventional diffusive transport theory. Finally, for MD simulations of GPs assembling in the absence of a membrane the slowdown of assembly rates as shells near completion at **ρ** = 2*R*_0_ is negligible.

**Fig 5.**
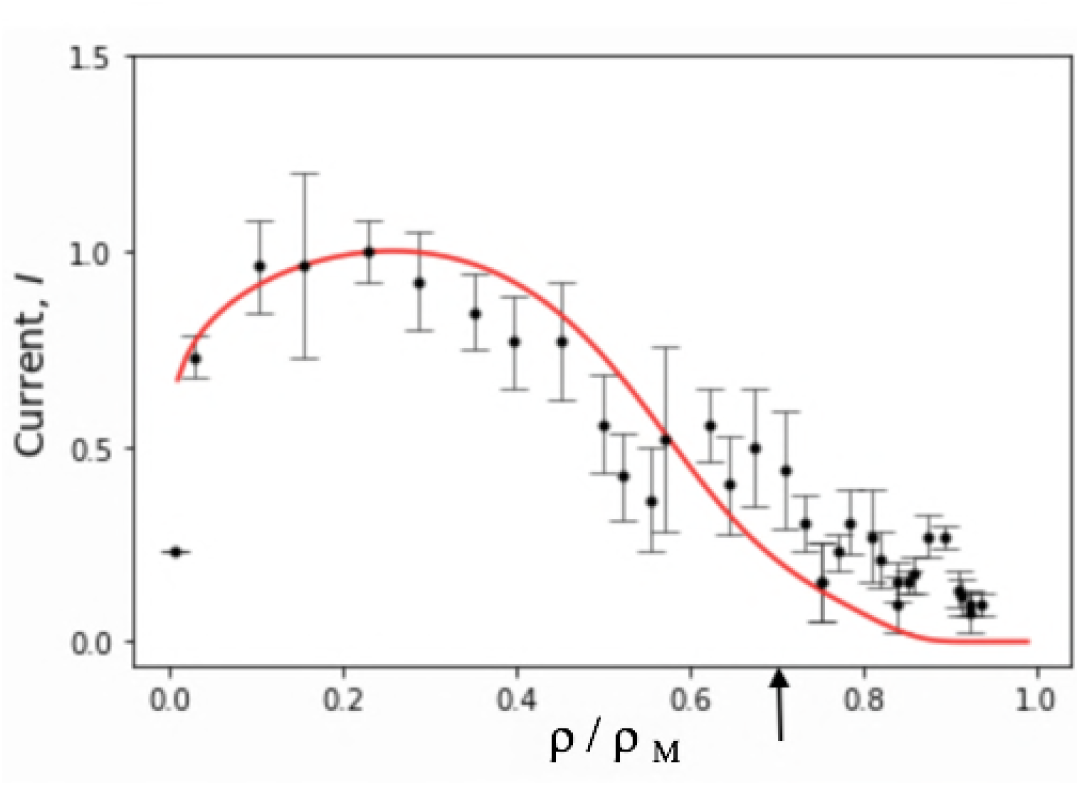
Solid red line: Diffusion current *I* from the exterior to the growing bud versus relative size **ρ**/**ρ*_M_* of the bud computed from Eq.7. *γ* = 0.9 was the sole fitting parameter. The current and the radius were normalized with respect to their maximum values. Black dots: assembly current obtained from the Brownian Dynamics simulation of Fig.2. Error bars were obtained by averaging over three runs. The strength *ϵ*_gg_ of the interaction between the capsid proteins was 6*k*_B_*T*. The black arrow indicates the location of the maximum of the current profile predicted by conventional diffusive transport theory (i.e. *γ* = 0 in Eq.11).

## Conclusions

In summary, the physics of diffusion of proteins along a curved surface provide a natural physical mechanism for the pausing and stalling observed during late-stage budding of enveloped viruses in general and of HIV-1 in particular. Capsid proteins diffusing in from infinity towards the growth surface must pass through a neck region with negative Gauss curvature. This requirement applies even to assembly processes in which RNA-protein interactions play a role [38] or proteins are targeted near capsid assembly sites [39]. The associated energy barrier does not show up in the equilibrium thermodynamics. The importance of the geometrical barrier effect is determined by the dimensionless parameter 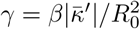. If *γ* is of the order of one or larger, then the suppression of the current by the geometrical barrier shows up already at relatively large aperture angles while for smaller *γ,* the effect appears only for increasingly smaller apertures. One can estimate 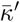 by the condition that 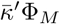 should be of the order of the Gauss curvature modulus 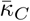 of the capsid. Here, Φ*_M_* ≃ *1*/*a*^2^ is the protein concentration of the assembled shell *a* is of the order of a nanometer. Numerical estimates of Gauss moduli [40] typically lead to 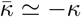 [41]. The curvature modulus of capsids is in the range of 100 *k_B_T* or more. It follows that 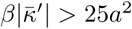, which means that *γ >* 25(*a*/*R*_0_)^2^. For a small enveloped virus, such as hepatitis D, the lower bound 25(*a*/*R*_0_)^2^ is in the range of 0.1 to 1.0. The dependence of the growth rate on the amount of capsid material is consistent with numerical simulations of the assembly of the alphavirus with values for *γ* in this range.

According to the numerical simulations, there is a critical point as a function of protein-protein interaction strength *ϵ*_gg_ below which assembly stalls and above which the assembly completes and scission is spontaneous. This provides an explanation of the observation that the protein shells of some enveloped viruses, like HIV-1, have a hole at the pinch-off -site while others - such as Herpes – have completely closed protein shells. Consistent with this explanation is the fact that the interaction strength between the capsid proteins of immature retroviruses is believed to be quite weak [42], which could place them below the critical point. Even when scission was spontaneous in the simulations, there was still a pronounced slow down of the assembly process prior to scission. This suggests that measurement of the budding kinetics of enveloped viruses that do not have a hole in their capsids, such as Herpes, still should encounter a clear slow down prior to scission.

The simulations show that scission is spontaneous for sufficiently large values of *ϵ*_gg_. Why do viruses recruit the ESCRT machinery instead of taking this route? A possible reason could be that, according to the numerical simulations, the capsid becomes increasingly *defected* for large *ϵ*_gg_ and this could interfere with other functions of the virus. There is another manner in which the geometrical barrier could be suppressed namely by decoupling the capsid proteins from the Gaussian curvature so 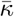 and 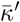 both are negligible. This may not be an option either because the spherical cap shape becomes – as we discussed – mechanically unstable when 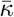 goes to zero. The opportunistic recruitment of the cell machinery – sometimes referred to as a “Trojan Horse” strategy [43]– avoids all these problems.

The geometrical barrier could be an interesting target for anti-viral strategies. If it would be possible to engineer “Gaussian proteins” with an affinity for lipid bilayers with negative Gauss curvature, then such proteins could stabilize the neck against attack by the ESCRT machinery proteins and halt the budding process.

## Materials and methods

Our simulations employ a coarse-grained model that was designed to capture the essential physical features of the membrane and alphavirus transmembrane glycoproteins (GPs) (see Fig. 6). Although the GPs are transmembrane proteins, their assembly is described by the same continuum model as HIV capsid proteins adsorbed to the membrane. Moreover, in our model the conical regions which drive curvature of the model subunit oligomers are located within and below the plane of the membrane, as we found that this arrangement facilitated completion of assembly [1]. We note that the stalling described in the main text was observed for all subunit interaction geometries that we have considered for the alphavirus model [1], as well as in another model for proteins that adsorb onto the membrane [44], suggesting that the barrier is a generic feature of assembly and budding on a membrane. We note that the stalling described in the main text was not observed in some previous budding simulations because they only considered early stages of budding [45–47]. While Refs. [48, 49] did consider the entire budding process, their model represented the capsid proteins as patchy spheres and the membrane as a triangulated monolayer, which likely eliminated or minimized coupling of the proteins to membrane Gaussian curvature.

**Fig 6.**
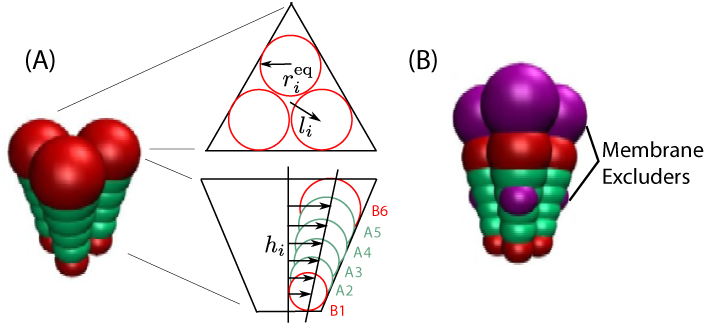
**(A)** (left) Image of a trimer subunit, with attractors (‘A_2_’-‘A_5_’) in green and excluders (‘B_1_’ and ‘B_6_’) in red. (right) Schematic of the subunit geometry, with views from directly above the plane of the membrane and within the plane of the membrane. Membrane excluders are not shown in these schematics to aid visual clarity. **(B)** Image of a subunit trimer, showing attractors (green, type ‘A’), excluders (red, type ‘B’), and membrane excluders (magenta, type ‘VX’).

We begin with an overview of each component of the model, and then give the full set of interaction potentials in section.

## Glycoproteins and capsid

Our model GPs were designed to roughly match the triangular shape, dimensions and aspect ratio of trimers-of-heterodimers of E1 and E2 GPs in the Sindbis virion [1, 50, 51]. There are 80 of these trimers arranged with T=4 icosahedral symmetry in the virion structure. On the capsid surface each trimer forms a roughly equilateral triangle with edge-length ~ 8nm. In the radial direction, each E1-E2 heterodimer spans the entire lipid membrane and the ectodomain spike, totaling ~ 12nm in length.

Our GP trimer subunit comprises three cones, which are fused together and simulated as a rigid body. Each cone is represented by an array of 6 beads of increasing diameter, following the model described by Chen et al. [52]. However, our cones are truncated, so that they form a shell with an empty interior, as shown in Fig. 6. The cones experience lateral interactions, with a preferred angle that, in the absence of a membrane, drives assembly into shells with a typical size of 80 trimers, with fluctuations in the range 79 – 82 trimers. The attractions are mediated by the four interior pseudoatoms within each cone (A2-A5 in Fig. 6), while the innermost and outermost pseudoatoms (B1 and B6) experience only excluded volume.

## Lipid membrane

The lipid membrane is represented by the implicit solvent model from Cooke and Deserno [13]. This model enables on computationally accessible timescales the formation and reshaping of bilayers with physical properties such as rigidity, fluidity, and diffusivity that can be tuned across the range of biologically relevant values. Each lipid is modeled by a linear polymer of three beads connected by FENE bonds; one bead accounts for the lipid head and two beads for the lipid tail. An attractive potential between the tail beads represents the hydrophobic forces that drive lipid self-assembly. For the simulations described here, the membrane bending modulus was set to *κ*_mem_ ≈ 12.5*k*_B_*T*.

## Glycoprotein-membrane interactions

We used a minimal model for the GP-membrane interaction. We add six membrane excluder beads (type ‘VX’) to our subunit, three at the top and three at the bottom of the subunit, with top and bottom beads separated by 7nm (magenta beads in Fig. 6B). These excluder beads interact through a repulsive Lennard-Jones potential with all membrane beads, whereas all the other cone beads do not interact with the membrane pseudoatoms. In a simulation, the subunits are initialized with membrane located between the top and bottom layer of excluders. The excluded volume interactions thus trap the subunits in the membrane throughout the length of the simulation, but allow them to tilt and diffuse laterally.

## GP Conformational changes and implementation of constant GP concentration

Experiments suggest that viral proteins from many families interconvert between ‘assembly-active’ and ‘assembly-inactive’ conformations, which are respectively compatible or incompatible with assembly into the virion [53–55]. Experiments suggest imilar conformational changes for the alphavirus GPs E1 and E2 [55, 56]. Computational modeling suggests that such conformational dynamics can suppress kinetic traps [57, 58]. Based on these considerations, our GP model includes interconversion between assembly-active and assembly-inactive conformations. The two conformations have identical geometries, but only assembly-active conformations experience attractive interactions to neighboring subunits. I.e., there are no attractive interactions (Eq. 18 below) for subunit pairs in which one or both of the subunits is in the inactive conformation.

We adopt the ‘Induced-Fit’ model of Ref. [57], meaning that interaction with an assembling GP shell or the NC favors the assembly-active conformation. For simplicity, we consider the limit of infinite activation energy. In particular, with a periodicity of *τ*_c_ all the inactive subunits found within a distance 1.0*σ* of the capsid are switched to the active conformation, while any active subunits further than this distance from an assembling shell convert to the inactive conformation. Results were unchanged when we performed simulations at finite activation energies larger than 4*k*_B_*T*.

To maintain a constant subunit concentration within the membrane (outside of the region where an assembling shell is located) we include a third subunit type called ‘reservoir subunits’, which effectively acts as a reservoir of inactive subunits. These subunits interact with membrane beads but experience no interactions with the other two types of GP subunits. With a periodicity of *τ*_c_, reservoir subunits located in a local region free of active or inactive subunits (corresponding to a circumference of 1.5 times the radius of the largest subunit bead) are switched to the assembly-inactive state.

## Subunit geometry

The geometry of the model GP trimer subunit is schematically shown in Fig. 6. As explained above, the subunit consists of three cones symmetrically placed around the subunit axis. Each cone contains six pseudoatoms. Only the inner four pseudoatoms (denoted as A) experience attrative interactions. The outer two pseudoatoms, B, interact with the rest through excluded volume. The pseudoatoms are placed at heights *h_i_* = [16.0, 17.5, 19.0, 20.5, 22.0, 23.5]*σ*. At each plane *z* = *h_i_* there are three identical pseudoatoms forming an equilateral triangle of radius *l_i_* = *h_i_* tan *α_l_,* where *α_l_* can be tuned. Since assembly in bulk is slightly more robust for smaller *α_l_,* we choose an optimal value *α_l_* = 7*°*. The radius of each pseudoatom is then given by *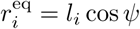*, with *ψψ* = 94.9° the parameter that controls the preferred curvature of the subunits. Finally, to embed the subunits in the membrane we add two layers of three membrane excluders ‘VX’, consistent with the cone geometry, at height *h*_in_ = 19.0*σ* (inner domain) and *h*_out_ = 26.0*σ* (outer domain). The sequence of pseudoatoms across the shell reads [B_1_,A_2_,A_3_,VX_in_,A_4_,A_5_,B_6_,VX_out_].

## Simulations

We performed simulations in HOOMD-blue [59], version 1.3.1. Both the subunits and the NC were simulated using the Brownian dynamics algorithm for rigid bodies. The membrane dynamics was integrated using the NPT algorithm, a modified implementation of the Martina-Tobias-Klein thermostat-barostat. The box size changes in the membrane plane, to allow membrane relaxation and maintain a constant lateral pressure. The out-of-plane dimension was fixed at 200*σ*.

Our simulations used a membrane patch with size 170 × 170nm^2^ (*A* ~ 28, 900nm^2^), which contained 51, 842 lipids. We compared membrane deformations, capsid size and organization from these simulations against a set of simulations on a larger membrane (210 × 210nm^2^, *A* ~ 44, 100nm^2^) and observed no significant differences, suggesting that finite size effects were minimal. Simulations were initialized with 160 subunits uniformly distributed on the membrane, including 4 active-binding subunits (located at the center of the membrane) with the remainder in the assembly-inactive conformation. In addition, there were 156 subunits in the reservoir conformation uniformly distributed.

The membrane was then equilibrated to relax any unphysical effects from subunit placement by integrating the dynamics for 1,500 *τ*_0_ without attractive interactions between GPs. Simulations were then performed for 4,200 *τ*_0_ with all interactions turned on. The timestep was set to Δ*t* = 0.0015, and the thermostat and barostat coupling constants were *τ_T_* = 0.4 and *τ_P_* = 0.5, respectively. Since the tension within the cell membrane during alphavirus budding is unknown, we set the reference pressure to *P_0_* = 0 to simulate a tensionless membrane. The conformational switching timescale was set to *τ_c_* = 3*τ*_0_, sufficiently frequent that the dynamics are insensitive to changes in this parameter.

## Acknowledgments

We would like to thank Joseph Rudnick for providing us with the solution of Eq.7, Markus Deserno for reading a first draft of the paper and the NSF-DMR for support under Grant 1610384 (SD, BS, RB); the Brandeis Center for Bioinspired Soft Materials, an NSF MRSEC, DMR-1420382 (GRL); and the NIH, Award Number R01GM108021 from the National Institute Of General Medical Sciences (GRL and MFH). Computational resources were provided by the NSF through XSEDE computing resources (MCB090163) and the Brandeis HPCC which is partially supported by the Brandeis MRSEC.

## Appendix Interaction potentials

### Interaction potentials

The total interaction energy *U*_tot_ can be separated into two contributions,

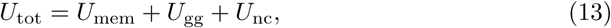

where *U*_mem_ represents the interaction energy between the membrane beads and *U*_gg_ accounts for the interaction of between subunits as well as with the membrane.

### Membrane interactions

The membrane lipids consist of three beads, the first representing the lipid head and the other two connected through two finite extensible nonlinear elastic (FENE) bonds with maximum length *r*_cut_ = 1.5*σ*,

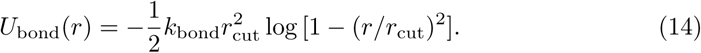

with *k*_bond_ = 30*ϵ*_0_/*σ*^2^. A harmonic spring links the two outer beads, to ensure that the lipids maintain a cylindrical shape,

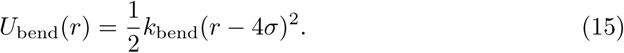

All membrane beads interact via a Weeks-Chandler-Andersen potential,

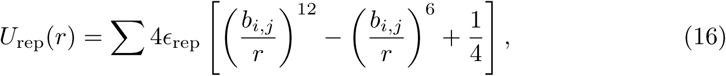

with *ϵ*_rep_ = 1 and cutoff *r*_cut_ = 2^1/6^*b_i,j_*. The parameter *b_i,j_* depends on the identities of the interacting beads: *b*_h,h_ = *b*_h,t_ = 0.95*σ* and *b*_t,t_ = 1.0*σ*, with the subscripts ‘h’ and ‘t’ denoting head and tail beads, respectively. The hydrophobic nature of the lipid tails is accounted for by an attractive interaction between all pairs of tail beads:

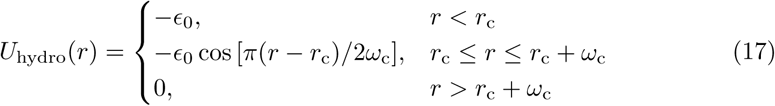

with *ϵ*_0_ = 1.0, *r*_c_ = 2^1^/^6^*σ*. The potential width *ω*_c_ is a control parameter that determines, among other properties, the membrane rigidity. Unless otherwise specified, *ω*_c_ = 1.6.

### GP-GP interactions

The interaction potential between pairs of GP subunits, *U*_gg_, consists of two terms. If both subunits are in the active conformation, there is an attractive interaction between pairs of attractor pseudoatoms ‘A’, modeled by a Morse potential. Beads interact only with those of the same kind on a neighboring cone, A*_i_*-A*_i_*, *i* = 2,..., 5, and the equilibrium distance of the potential depends on the pseudoatom radius, 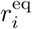:

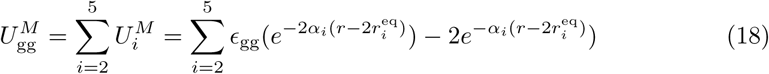

With *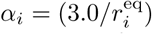*. The cutoff of this interaction was set at *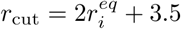*. All subunit beads of type ‘A’ and ‘B’ experience excluded volume interactions regardless of whether subunits are in the active or inactive conformations, according to:

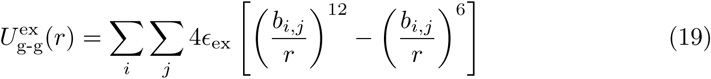

with *ϵ*_ex_ = 1.0 and cutoff radius 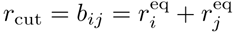. The sum extends to all the subunit beads of type ‘A’ and ‘B’.

In the subunits, only the pseudoatoms ‘VX’ interact with the membrane beads; there is no interaction between membrane beads and ‘A’ or ‘B’ pseudoatoms. The interaction between subunit excluders and membrane beads corresponds to the repulsive part of the Lennard-Jones potential,

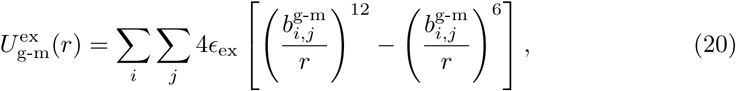

where *i* runs over all lipid beads and *j* over all ‘VX’ pseudoatoms, and 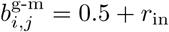 for the inner excluders VX_in_ and *b_i,j_* = 0.5 + *r*_in_ for the outer excluders VX_out_.

### Modulus values

The mean curvature modulus for this model was calculated in Ref. [1] to be *κG* ≈ 25.66*ϵ*_gg_, and the calculation in Ref. [41] for a related model shows that *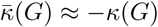.* However, there is an additional energetic penalty (not present in the Helfrich hamiltonian) for regions in which the two principle curvature are mismatched, so the continuum description of the bending energy goes as [41].

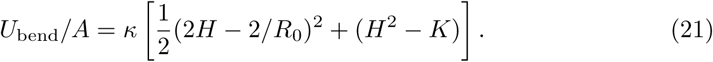

## Footnotes

1 This is ultimately a consequence of the Theorema Egregium of Gauss.

